# An automated workflow for multiplexed single-cell proteomics sample preparation at unprecedented sensitivity

**DOI:** 10.1101/2021.04.14.439828

**Authors:** Claudia Ctortecka, David Hartlmayr, Anjali Seth, Sasha Mendjan, Guilhem Tourniaire, Karl Mechtler

**Affiliations:** Research Institute of Molecular Pathology (IMP), Vienna BioCenter (VBC), Campus-Vienna-Biocenter 1, 1030 Vienna, Austria; Cellenion SASU, 60F avenue Rockefeller, 69008 Lyon, France; Institute of Molecular Biotechnology of the Austrian Academy of Sciences (IMBA), Vienna BioCenter (VBC), Dr. Bohr-Gasse 3, 1030 Vienna, Austria; The Gregor Mendel Institute of Molecular Plant Biology of the Austrian Academy of Sciences (GMI), Vienna BioCenter (VBC), Dr. Bohr-Gasse 3, 1030 Vienna, Austria

## Abstract

The analysis of single-cell proteomes has recently become a viable complement to transcriptomics and genomics studies. Proteins are the primary driver of cellular functionality and mRNA levels are often an unreliable proxy of such. Therefore, the global analysis of the proteome is essential to study cellular identities. Multiplexed and label-free mass spectrometry-based approaches with single-cell resolution have lately attributed surprising heterogeneity to presumed homogenous cell populations. Even though specialized experimental designs and instrumentation have demonstrated remarkable advances, the efficient sample preparation of single cells still lag. Here, we introduce the proteoCHIP, a universal option for single-cell proteomics sample preparation at surprising sensitivity and throughput. The automated processing using a commercial system combining single-cell isolation and picoliter dispensing, the cellenONE^®^, reduces final sample volumes to low nanoliters submerged in a hexadecane layer simultaneously eliminating error-prone manual sample handling and overcoming evaporation. The specialized proteoCHIP design allows direct injection of single cells via a standard autosampler resulting in around 1,500 protein groups per analytical run at remarkable reporter ion signal to noise while reducing or eliminating the carrier proteome. We identified close to 2,600 proteins across 170 multiplexed single cells from two highly similar human cell types. This dedicated loss-less workflow allows distinguishing *in vitro* co-differentiated cell types of self-organizing cardiac organoids based on indicative markers across 150 single cells. In-depth characterization revealed enhanced cellular motility of cardiac endothelial cells and sarcomere organization in cardiomyocytes. Our versatile and automated sample preparation has not only proven to be easily adaptable but is also sufficiently sensitive to drive biological applications of single-cell proteomics.

## Introduction

In recent years, single-cell analysis has demonstrated valuable insights into heterogeneous populations. However, proteins, especially their post translational modifications, are the primary driver of cellular identity and function. Therefore, complementing single-cell transcriptomics and genomics approaches with global protein analysis is essential. However, most protein profiling techniques with single-cell resolution, still rely on the availability of antibodies. Continuous technological advances drive sensitivity and accuracy of mass spectrometry (MS)-based single-cell proteomics (SCP) for hypothesis-free measurements. Despite that, three main aspects, throughput, measurement variability, and, most importantly, sample preparation efficiency still lag behind comparable sequencing techniques.

The combination of dedicated instrumentation with sensitive acquisition strategies for label-free single-cell analysis has been demonstrated highly accurate but very limited in throughput.^1,2^ In label-free MS experiments single cells are processed and analyzed individually, prone to peptide losses and poor throughput. This was addressed through isobaric labeling (i.e. tandem mass tags – TMT), allowing to uniquely barcode individual cells for simultaneous analysis and relative quantification upon MS-based investigation.^3^ TMT reagents are available in several multiplexing capacities enabling the analysis of up to eighteen samples in one experiment.^4^ This not only reduces the measurement time per single cell from about 25 minutes for label-free analysis to merely 3.5 minutes but also increases the input material per sample. For multiplexed experiments, single cells are processed individually, TMT-labeled after tryptic digestion and combined into one sample for MS analysis. The identical mass of the TMT labels allows for the simultaneous elution of a peptide from all samples, therefore multiplying the precursor signal and contributing ions for peptide identification. Moreover, the differently equipped heavy isotopes of the TMT-reagents enable relative quantification upon fragmentation in the MS. This was previously adopted to combine single cells with an abundant carrier spike for improved precursor selection, increased fragment ions for identification and reduced peptide losses throughout the workflow (SCoPE-MS).^5^ However, such extreme carrier ratios were demonstrated to impair quantitative accuracy and possibly affect biological conclusions.^6,7^

Similarly, peptide losses during sample preparation were minimized with dedicated slide-based SCP sample preparation workflows, such as the nanoPOTS (nanodroplet processing in one-pot for trace samples) or nPOP.^1,8^ These droplet-based sample preparations minimize sample volumes to less than 100 nL, which were automated via the cellenONE^®^, a commercial liquid dispensing instrument for cell isolation and single-cell preparation.^8–11^ The latest generation nested nanoPOTS and nPOP, smartly unify TMT sets with a microliter droplet on top of each sample array, extensively reducing manual handling. Nevertheless, to boost identifications they combine the single cells with an abundant >10 ng bulk carrier and manually transfer the combined sample to a HPLC vial or rely on a custom-built nanoPOTS autosampler.^11,12^ To address those shortcomings, we here describe a highly versatile and automated workflow for isobarically multiplexed single-cell proteomics sample preparation at unprecedented sensitivity. We present the proteoCHIP for cross-contamination-free processing of up to 592 single cells per experiment. Moreover, our dedicated proteoCHIP design overcomes all manual sample handling and directly interfaces with standard autosamplers for accurate quantification and comparable protein identifications.

## Main

### Multiplexed single-cell proteomics sample preparation with the proteoCHIP

We here introduce the proteoCHIP as a viable option for automated single-cell proteomics sample preparation within a robotic platform combining single-cell isolation and picolitre dispensing, the cellenONE^®^. The proteoCHIP nanowell part is designed to process 192 single-cells simultaneously (12 multiplexed sets of 16 samples) and up to 576 single cells per experiment (i.e. three proteoCHIPs; Fig. 1a-b). The PTFE-based chip in the size of a standard microscopy slide overcomes peptide losses to plastic or glass ware with similar GRAVY indices compared to dedicated low-binding and PEG-coated autosampler vials (Supplemental Fig. 1).^13,14^ After automatic pooling via a benchtop centrifuge to the proteoCHIP funnel part, that is directly interfaced with a standard HPLC autosampler for improved peptide recovery (Fig. 1c). Moreover, we overcome evaporation of low nanoliter sample volumes with an oil layer for constant enzyme and chemical concentration. To not interfere with HPLC injection, we chose hexadecane, which solidifies at autosampler temperature. Despite aiming to minimize reaction volumes, the proteoCHIP nanowells hold up to 600 nL allowing to readily adapt the protocol without cross-contamination of the samples.

**Figure 1:**
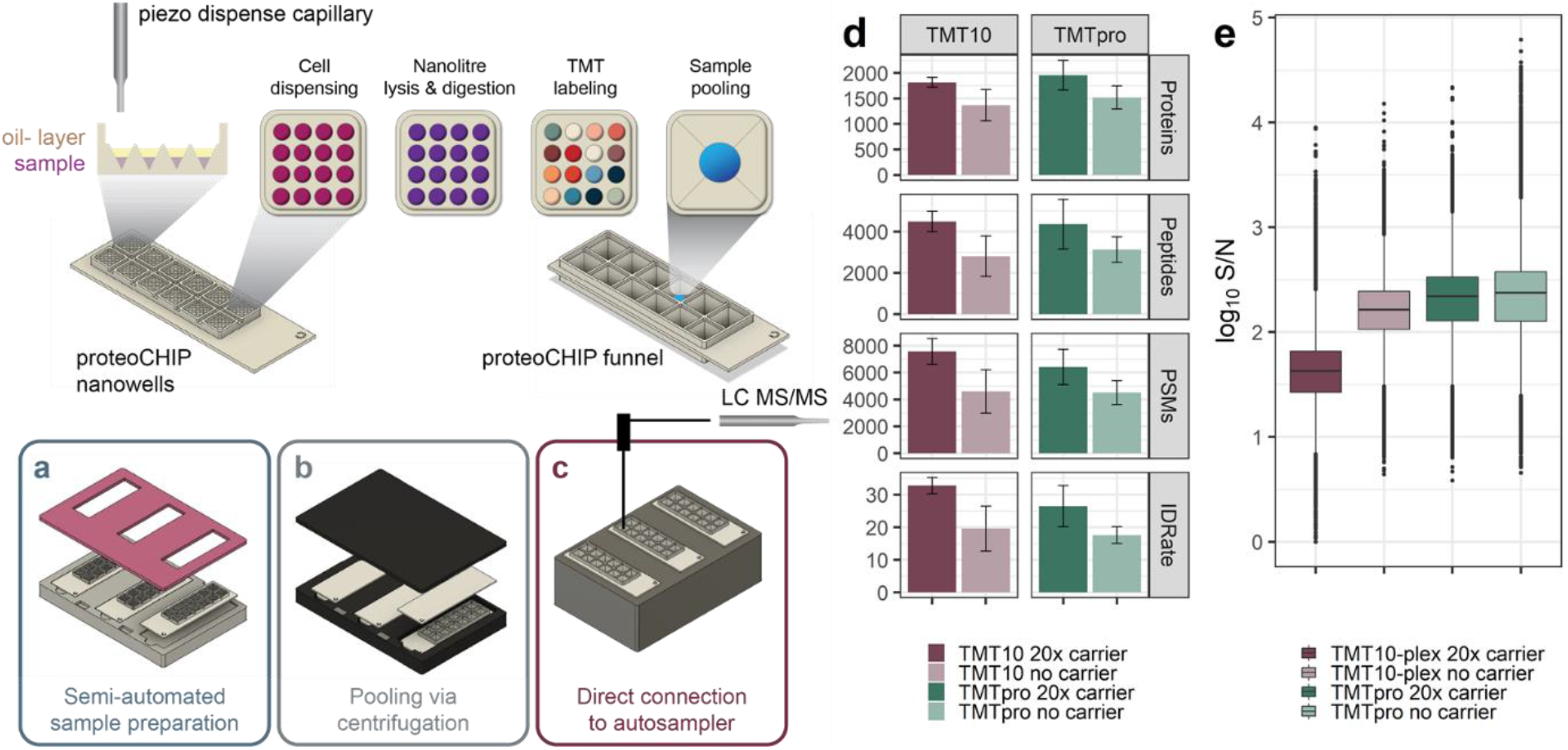
The proteoCHIP multiplexing workflow for TMT10 and TMTpro. **(a)** Up to sixteen nanowells/single-cells per TMT set are prepared inside the cellenONE^®^, **(b)** automatically combined via benchtop centrifugation and **(c)** directly interfaced with a standard autosampler for loss-less acquisition. **(d)** Identified proteins, peptides, PSMs, ID-rate and **(e)** RI S/N of TMT10 (red) or TMTpro (green) samples with 20x or no carrier.

Based on the loss-less proteoCHIP design, we speculate that an abundant carrier is not required to generate protein IDs comparable to other techniques.^11,15^ For this, we sorted 100 or 20 cells into one nanowell per single-cell sample set. While the 20x carrier elevated protein IDs by 24%, increasing the carrier ratio to 100x improves protein IDs by 42% (Supplemental Fig. 2a). Nevertheless, close inspection revealed drastic reporter ion (RI) signal suppression of over 40% by the 20x carrier and additional 25% decrease by the 100x carrier spike (Supplemental Fig. 2b). As high RI signal to noise (S/N) parallels with improved quantitative accuracy we conclude that the carrier not only impairs quantitation but also data completeness and expedites missing RI frequency (Supplemental Fig. 2b-c).^6,7^ Next, we aimed at determining if precursor selection is influenced by the carrier ratio or preparation. For this, we spiked bulk digested equivalents of 20 or 100 cells, sorted 20 cells and no carrier. As expected, the 20x sorted carrier increased protein IDs by ∼30%, however, the equivalent bulk diluted carrier merely elevated identified proteins, even decreasing peptides by 10% in comparison to the no-carrier sample (Fig. 1d; Supplemental Fig. 2a, d). Moreover, this effect was aggravated by increasing the carrier spike to 100x with similar protein IDs but 35% less PSMs (Supplemental Fig. 2d). Interestingly, an intersection of unique peptide sequences revealed distinct sets for each sample, largely paralleling with total peptides identified. However, from the 100x bulk carrier, despite yielding the lowest absolute number of peptide sequences over 20% were unique to this sample set (Supplemental Fig. 2e). We speculate that this is due to the dissimilar digestion conditions, thus highlighting carrier rather than single-cell precursors. We therefore conclude that a maximum of 20 sorted carrier cells suffices to boost ions but retain quantitative quality, limit missing data and independent precursor selection (Supplemental Fig. 2a-e).

Based on this, we processed multiple SCP batches with both available TMT tags, the TMT10 and TMTpro, with and without a 20x sorted carrier to compare reagents at optimized conditions for loss-less proteoCHIP sample preparation. We readily identify on average 2,071 and 1,539 protein groups based on 4,602 and 2,871 peptides per TMT10 set with a 20x or no carrier, respectively (Fig. 1d). All TMT10 single-cells combined (i.e. 306 single-cells) yield 2,913 protein groups based on 11,791 peptides. Similarly, the 20x and no carrier TMTpro samples (i.e. 276 single-cells) result in an average of 1,940 and 1,598 protein groups from 4,411 and 3,588 peptides per analytical run, respectively (Fig. 1d). Across all 276 TMTpro labeled single cell sets we identify over 3,674 protein groups from 14,830 peptides. Interestingly, TMTpro experiments with and without the carrier resulted in comparable protein identifications but higher S/N of the single-cell channels compared to the TMT10 (Fig. 1e).^7^ This is presumably due to the different fragmentation behavior of the longer TMTpro tag.^4^ Nevertheless, single cell RI S/N is reduced by 74% or 8% in the 20x versus the no-carrier samples, for TMT10 and TMTpro, respectively (Fig. 1e). Despite the low ratio between the carrier to single-cells, we speculate that the larger proportion of ions sampled from the carrier compresses the single-cell RI signal into the noise (Fig. 1e).^6,7^ Although the carrier undoubtedly boosts identifications in SCP, the reduced RI S/N and precursor selection influence suggests to keep the carrier ratio to a minimum (Fig. 1d-e; Supplemental Fig. 2a-e).^6,7,10^

For quantitative reproducibility and data quality evaluation, we compared missing quantitative signal per PSM and cumulative missing data across multiple analytical runs. For both labeling reagents, with or without a carrier, the RI S/N of two individual sample channels correlates well (i.e., r = >0.85) and outperforms previous reports of multiplexed SCP (Fig. 2a-d).^11,15^ This results in over 90% of all PSMs with maximum of one quantitative channel missing across all experimental setups (Fig. 2e-h). However, despite the low missingness per analytical run, precursor stochasticity accumulates distinct sets of identified precursors, thus decreasing the sample overlap (Fig. 2i-l). We speculate that the TMT labels and other reagents contribute to background signal and chemical noise, which interferes with precursor selection and MS/MS identification. Data aggregation of three analytical runs for both reagents (i.e. 30-48 single cells) reduces common protein identifications by 50% without matching between runs (Fig. 2i-l), as previously reported by others.^15,16^ Despite outstanding data quality, standard data dependent acquisition (DDA) accumulates missing data in large sample cohorts resulting in limited replicate overlap and requiring computational generation of quantitative values, which has to be considered prior to any SCP experiment.^17,18^

**Figure 2:**
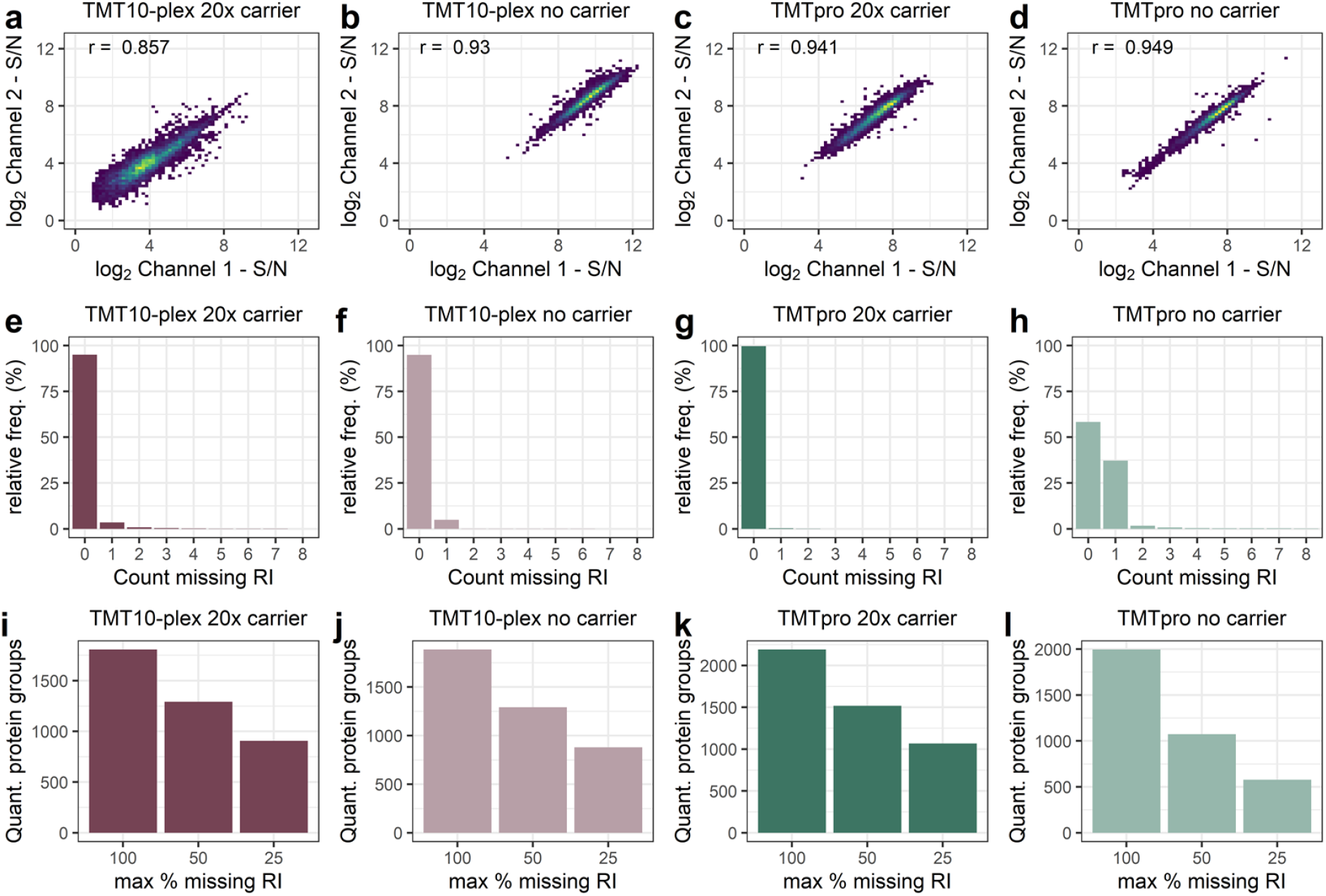
Data completeness and reproducibility of multiplexed single-cell proteomes with TMT10 and TMTpro reagents at different carrier ratios. Log2 S/N correlation of two single-cell samples for **(a)** TMT10 20x or **(b)** no carrier, **(c)** TMTpro 20x or **(d)** no carrier. r = Pearson correlation estimate. Percentage of relative missing RIs across three analytical runs per PSM for **(e)** TMT10 20x or **(f)** no carrier, **(g)** TMTpro 20x or **(h)** no-carrier samples. Cumulative missing RIs per quantified protein across three analytical runs for **(i)** TMT10 20x or **(j)** no carrier, **(k)** TMTpro 20x or **(l)** no-carrier samples.

### Cell type specific clustering of two human cell lines based on their single-cell proteome

Next, to benchmark our proteoCHIP-based sample preparation we aimed at differentiating two similarly sized human cell types based on their proteome (Supplementary Fig. 3a). For this we TMT10 labeled HeLa and HEK-293 cells without or with a 20x mixed carrier (50% HeLa and 50% HEK-293 cells). Across all 218 single cells we identified 2,598 proteins and 9,375 peptides with highly similar RI intensities for both cell types (Supplementary Fig. 3b). This not only indicates reproducible sample preparation but also strengthens our confidence that the two cell lines differ in proteome constitution rather than sample input. In detail, the 20x or no-carrier samples on average yield 1,812 or 1,477 proteins based on 4,517 or 3,348 peptides per analytical run (Fig. 3a). Similar to the HeLa-only samples a 20x carrier compressed the RI S/N by 57% while protein IDs only increased by 19% (Fig. 3a-b). Moreover, both the 20x and no-carrier samples demonstrated comparable replicate overlap, thus we continued without the carrier for enhanced quantitative data quality (Fig. 3c; Supplemental Fig. 3c).

**Figure 3:**
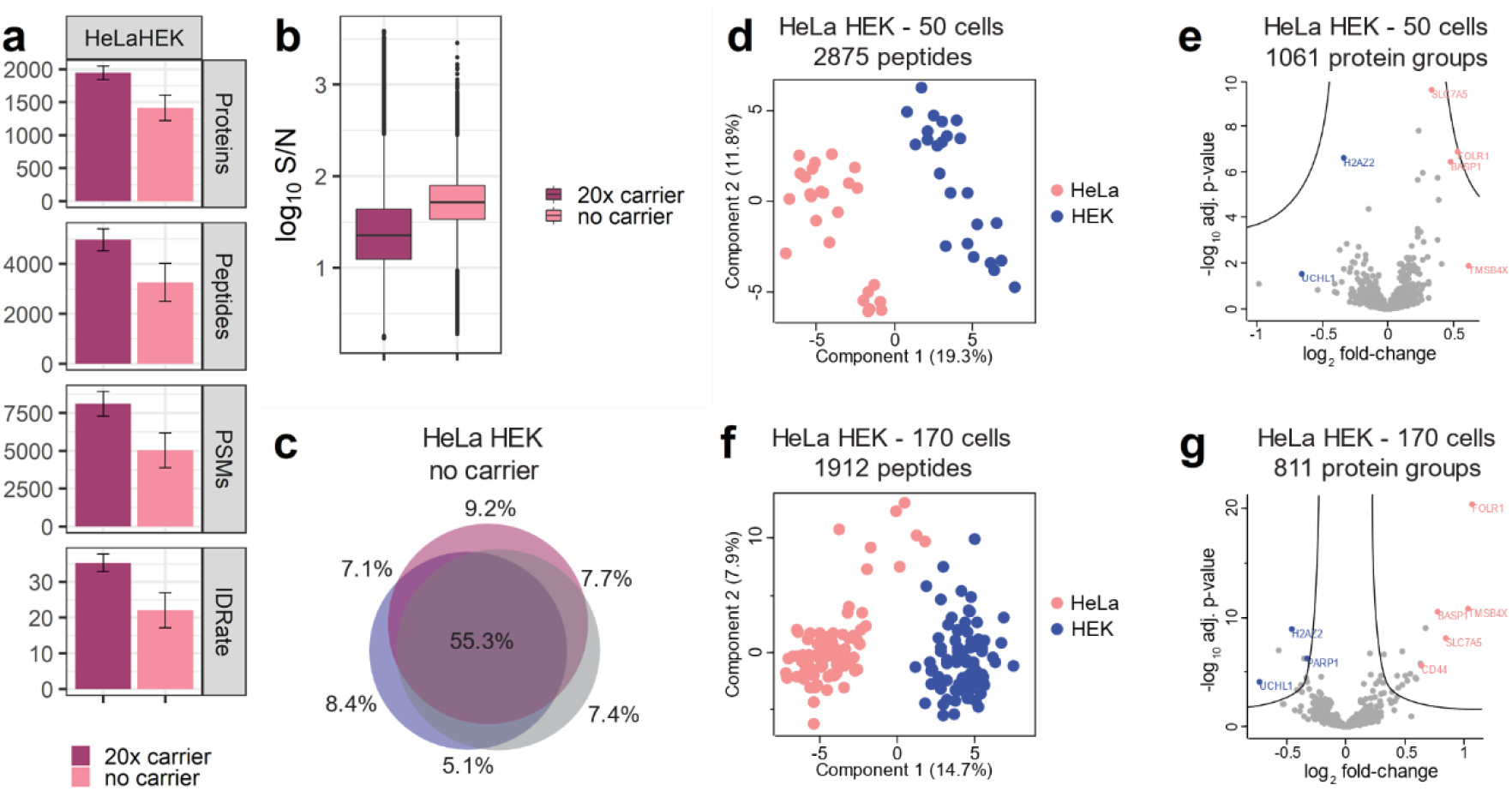
Comparison of HeLa and HEK-293 single-cell proteomes. **(a)** Protein groups, peptides, PSMs, MS/MS scans, ID-rate and **(b)** RI S/N of HeLa/HEK-293 samples with 20x and no-carrier. **(c)** Unique peptide sequence overlap across three analytical runs of HeLa/HEK-293 no-carrier samples. 50 single-cell **(d)** PCA and **(e)** differential protein abundance or 170 single-cell **(f)** PCA and **(g)** distinct proteins of HeLa (pink) and HEK-293 (blue). For volcano plots log2 fold change and −log10 adjusted p-value is shown.

We subjected 50 no-carrier HeLa/HEK-293 cells to principal component analysis (PCA) and observed distinct cell type specific separation via the first two components (Fig. 3d). Based on this, we assigned the respective sample channels to the two cell types and evaluated the top differentially expressed proteins (Fig. 3d-e). Interestingly, one of the top hits in HeLa cells is the brain acid soluble protein 1 (BASP1), which is downregulated in most tumor cell lines except some cervical cancer lines (Fig. 3e). In contrast to other cancer cell lines, the elevated levels of the tumor suppressor BASP1 in HeLa cells even promote tumor growth.^19^ Alongside BASP1, divergent proteins within our dataset parallel with normalized expression data obtained from the Human Protein Atlas^20^ (http://www.proteinatlas.org) (i.e. FOLR1, H2AZ2, SLC7A5, UCHL1, TMSB4X; Fig. 3e).

As many SCP applications require a more extensive sample cohort, we aggregated 170 HeLa/HEK-293 cells. The single cells intersect based on 1,912 unique peptide sequences with at least 70% data completeness and no match between runs or imputation (Fig. 3f). Despite rigorous quality controlling and minimal computational data generation, cell type cluster density decreases with sample size (Fig. 3d, f). We speculate that this is due to the reduced sample overlap through stochastic precursor selection and the thereof reduced analysis depth (Fig. 3c; Supplemental Fig. 3c-d). Nevertheless, we again assigned cellular identity to the samples within the two distinct clusters and subsequent inspection of the respective proteome characteristics corresponds to the 50-cell dataset and the Human Protein Atlas (Fig. 3e, g).^20^ We are therefore confident to accurately distinguish and identify cell types based on quantitative proteome differences across 170 single-cells (Fig. 3 d-g).

### Cell type-specific single-cell clustering of *in vitro* derived cardiac organoids

To assess our proteoCHIP workflow on a more tissue-like and biologically relevant sample, we characterized an *in vitro* derived cardiac organoid (cardioid).^21^ Cardioids are differentiated from human pluripotent stem cells using a chemically-defined protocol and develop into a cardiomyocyte organoid pervaded by a luminal structure lined with endothelial cells. These two cell types originate from a homogenous cell population within 7 days of mesoderm induction and can be visually distinguished by MYL7-GFP expression (i.e. Myosin regulatory light chain 7 indicative for cardiomyocytes). This not only allows to visually control for successful differentiation and estimation of relative cell proportions but also defines the ground truth before SCP analysis. Conveniently, the cellenONE^®^ ‘s ability to select cells based on their fluorescence signal allowed us to generate cardiomyocyte only (70 cells) and dual cell type (150 cells) single-cell samples. The 270 TMT10 labeled single cells yield 1,862 proteins based on 6,473 peptides, with 1,255 proteins on average for the cardiomyocytes only or 969 for the dual cell type samples (Fig. 4a). Despite low inter-sample protein identification variation, we speculate that proteome diversity and thus increased precursor pool complexity gives rise to the reduced replicate overlap and the decreased protein identifications of the dual versus the single cell-type sample (Fig. 1a; Supplemental Fig. 4a-c).

**Figure 4:**
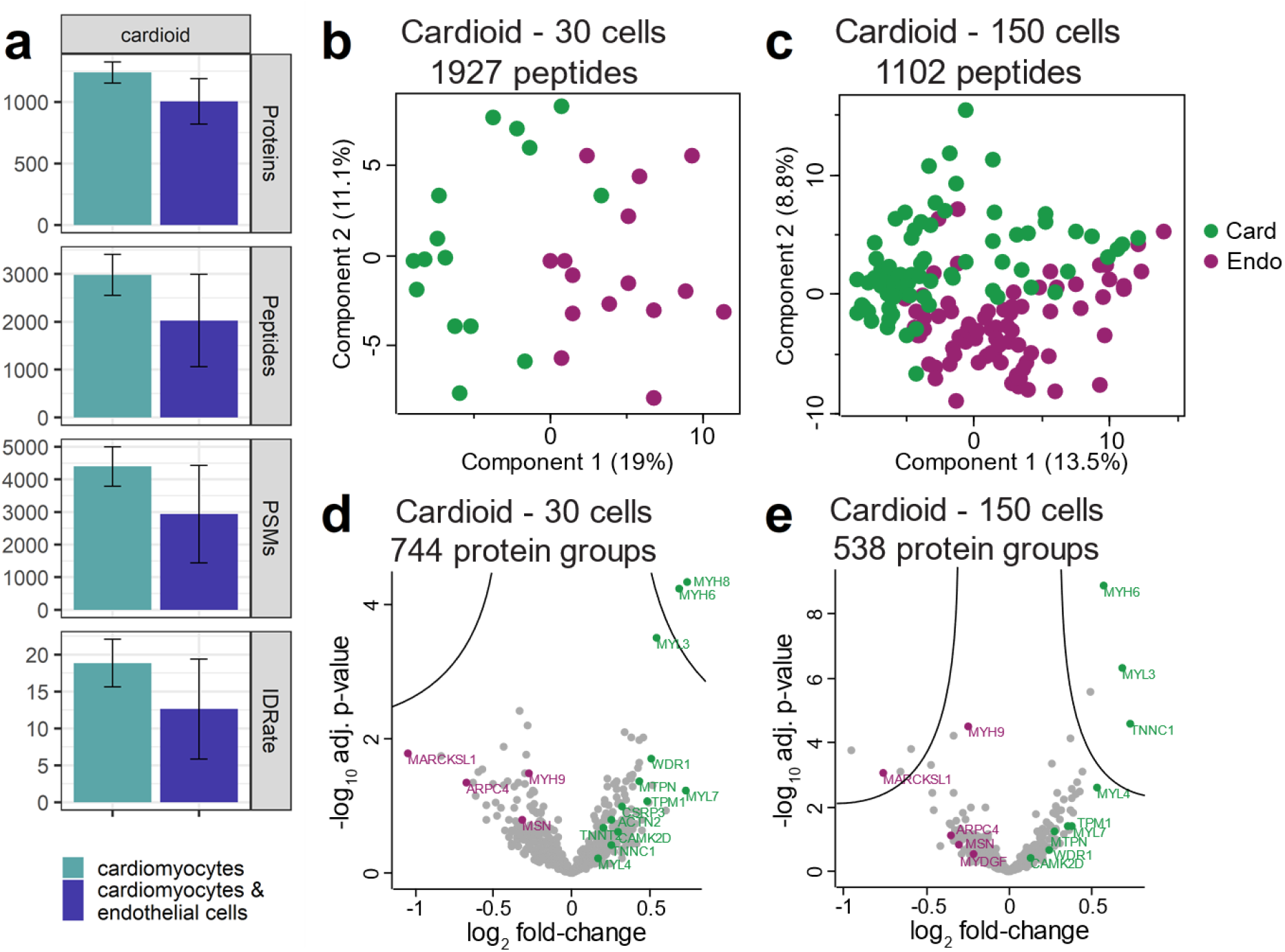
Cell type-specific separation of *in vitro* generated cardiac organoids. **(a)** Protein groups, peptides, PSMs, MS/MS scans and ID-rate of cardiac organoids. Single-cell proteome PCA of **(b)** 30 or **(c)** 150 single cells and proteome differences between cardiomyocytes (card - green) and endothelial cells (endo - purple) for **(d)** 30 or **(e)** 150 cells, respectively. For volcano plots, log2 fold change and - log10 adjusted p-value is shown.

Unbiased analysis based on 1,927 peptides across 30 dual cell-type cardioid single cells revealed that a comparable separation to the HeLa and HEK-293 samples without prior knowledge was not possible via PCA (Fig. 3d, f; Fig. 4b). We hypothesized that the co-differentiated cell types might vary in their proteome abundance and hence not constitute a clear two-cell type separation. We, therefore, increased the sample cohort 2 or 5-fold and observed a condensation of two partially converging clusters (Fig. 4b-c; Supplemental Fig. 4d). Close inspection revealed strong enrichment of indicative markers within the condensed populations. One of the most prominent marker for endothelial cells, MYH9 was highly enriched in the respective samples (Fig. 4d-e; Supplemental Fig. 4e).^20^ Interestingly, across the larger cohorts we additionally identified MYDGF in endothelial cells, which is described to stimulate endothelial cell proliferation, angiogenesis and inhibit cardiomyocyte apoptosis in a PI3K/AKT-dependent manner.^22^ Based on this we further examined our single-cell dataset for proteins involved in cellular motility and found ARPC4 and MARCKSL1, two actin regulators, to be significantly enriched in our endothelial cells. This indicates that the endothelial lining of the cardioids is tightly controlled and still actively undergoes rearrangement at day 7 post-induction.^23^ Contrarily, cardiomyocytes showed elevated levels of sarcomeric structural proteins MYL3, MYH6, TNNT2, TNNC1 and TNNI1 involved in muscle contraction.^20,21^ Apart from well-known marker we also identified WDR1, an actin interacting protein, to be highly abundant in cardiomyocytes. ^24,25^ The essential and highly conserved role of WDR1 in embryonic heart development and myocardial maintenance highlights the potential to study human cardiac organogenesis via SCP with *in vitro* cardioids.^21,26–28^ Moreover, our proteoCHIP dataset presents the first proteome characterization of cell type specific expression patterns and cellular dynamics in a developing human tissue at single cell resolution.

## Discussion

We here demonstrate the automation of SCP sample preparation using the proteoCHIP in conjunction with a commercial single cell isolation and picoliter dispenser, the cellenONE^®^. Our optimized protocol drastically reduces the digest volumes compared to previously published well-based techniques and is comparable to those successfully applied on modified glass slides.^8,11,15,29^ This not only minimizes chemical noise but the oil layer eliminates evaporation for constant enzyme and chemical concentrations. Next to unprecedented sample preparation efficiency, the specialized design of the proteoCHIP allows for automatic pooling of multiplexed samples to the proteoCHIP funnel using a standard benchtop centrifuge. This enables direct interfacing of the single cell samples with a standard autosampler for LC-MS/MS analysis without transfer nor specialized equipment (Fig. 1a-c).

Our semi-automated sample preparation eliminates all manual sample handling resulting in comparable protein identifications and drastically enhanced S/N of single cell RIs even at reduced or eliminated carrier (Fig. 1d-e; Supplemental Fig. 2a-c).^15,16^ This increases throughput by labelling single cells with all available TMT reagents and also improves the confidence of identifying peptides originating from single cells rather than the carrier (Supplemental Fig. 2d-e).^6^ Despite good correlation between individual samples (r = >0.85; Fig. 2a-d) and exceptional data completeness of individual samples (Fig. 2e-h) the identification overlap between analytical runs is still subject to improvement (Fig. 2i-l). Nevertheless, we successfully benchmark our proteoCHIP workflow for cell type-specific clustering of two human cell types with paralleling protein abundances to the Human Proteome Atlas for both cell types (Fig. 3d-g).^20^ Within this dataset we observed that precursor stochasticity in such low abundant samples results in sufficient sampling of abundant peptides but lower abundant precursors are often not shared across a large cohort. Our stringent quality metrics therefore drastically reduce the quantitative data and ultimately result in coalescent cell clusters (Fig. 3d, f; Supplemental Fig. 3d).

Moreover, our loss-less proteoCHIP sample preparation allowed us to characterize *in vitro* differentiated cardiac organoids resembling early human cardiac organogenesis. In contrast to the HeLa/HEK-293 samples, these yet immature and more similar cell types gave rise to converged clusters, which improved with increased sample cohort (Fig. 4b-c; Supplemental Fig. 4d). This indicated that the variance between individual cardioid cells and thus resulting background is higher compared to standard human cell lines and finally suggests more diverse sub-populations than initially expected (Fig. 4c). Remarkably, aside from the commonly known markers for cardiomyocytes and endothelial cells, we find strong elevation of the actin-binding/interacting proteins ARPC4 and MARCKSL1 (Fig. 4d-e; Supplemental Fig. 4e). This confirms active cellular migration of endothelial cells and indicates tightly controlled restructuring of the endothelial lining and possible early blood vessel formation.^23^ Similarly, we identify WDR1 to be highly expressed in cardiomyocytes, which has been sparsely connected to early cardiac development in mice and will be subject of further investigation utilizing our tightly controlled system.^21,26^ These small differences in protein abundances between cell types are likely to be unrecognized in bulk samples and therefore strengthen the importance of single-cell or subpopulation analysis.

In conclusion, to the best of our knowledge this is the first study successfully identifying and characterizing *in vitro* derived cell types from organoids through multiple markers across >100 single cells. We are therefore convinced that our proteoCHIP sample preparation in conjunction with dedicated acquisition strategies will increase analysis depth to profile such similar sub-populations based on their proteome. Moreover, the unprecedented RI S/N across multiple samples and >75% data completeness per analytical run with present acquisition strategies leaves us confident that multiplexed DIA or more selective DDA workflows will further advance data quality.^18,30^ The conjunction of our proteoCHIP with the cellenONE^®^ achieves efficient single-cell proteomics sample preparation, which can be readily adapted and overcomes detrimental peptide losses to in-depth characterize diverse samples.

## Material and Methods

### Sample preparation

HeLa and HEK293T cells were cultured at 37 °C and 5% CO2 in Dulbecco ‘s Modified Eagle ‘s Medium supplemented with 10% FBS and 1x penicillin-streptomycin (P0781-100ML, Sigma-Aldrich, Israel) and L-Glut (25030-024, Thermo Scientific, Germany). After trypsinization (0.05% Trypsin-EDTA 1x, 25300-054, Sigma-Aldrich, USA/Germany), cells were pelleted, washed 3x with phosphate-buffered saline (PBS) and directly used for single-cell experiments. Cardioids were cultured as previously described, briefly, WTC-MYL7-GFP cells were seeded in ultra-low attachment plates, after 24hrs cell aggregates were induced with CDM medium containing FGF2, LY294002, Activin A, BMP4, CHIR99021 and insulin. Cardiac differentiation was performed with CDM medium including BMP4, FGF2, insulin, IWP2 and retinoic acid for four days and CDM medium with BMP4, FGF2 and insulin for two days. Subsequently, cardioids were maintained in CDM with insulin.^21,31^ Samples were collected 7 days post induction and dissociated using the STEMdiff cardiomyocyte dissociation kit (#05025, Stem Cell Technologies) and subsequently washed 3x with PBS.

40 nL lysis buffer (0.2% DDM (D4641-500MG, Sigma-Aldrich, USA/Germany), 100 mM TEAB (17902-500ML, Fluka Analytical, Switzerland), 10 ng/uL trypsin (Promega Gold, V5280, Promega, USA) was dispensed into each well using the cellenONE^®^ (Cellenion, France) at elevated humidity. After single cell deposition (gated for cell diameter min. 22 μm and max. 33 μm, circularity 1.1, elongation 1.84) submerged with a layer of Hexadecane (H6703-100ML, Sigma-Aldrich, USA/Germany). The chip was incubated at 50 °C for 2 hours, directly on the heating deck inside the cellenONE^®^. For TMT multiplexing 60 nL of 100 mM TEAB and 100 nL of ∼10 mM TMT10 or TMTpro in anhydrous acetonitrile was added to the respective wells and incubated for 1 hour at room-temperature. TMT was subsequently quenched with 50 nL 0.5 % hydroxylamine (90115, Thermo Scientific, Germany) and 3% hydrochloric acid followed by sample pooling via centrifugation at 1,500 rpm for 2 minutes to the proteoCHIP funnel part for direct injection or transferred to 0.2 mL PCR-tubes coated with 1e-3 % Poly(ethylene glycol) (95172-250G-F, Sigma-Aldrich, Germany).

### LC-MS/MS analysis

Samples were measured on an Orbitrap Exploris™ 480 Mass Spectrometer (Thermo Fisher Scientific) with a reversed phase Thermo Fisher Scientific Ultimate 3000 RLSC-nano high-performance liquid chromatography (HPLC) system coupled via a Nanospray Flex ion source equipped with FAIMS (operated at −50 CV). Labeled peptides were first trapped on an Acclaim™ PepMap™ 100 C18 precolumn (5 μM, 0.3 mm X 5 mm, Thermo Fisher Scientific) and eluted to the analytical column nanoEase M/Z Peptide BEH C18 Column (130Å, 1.7 μm, 75 μm X 150 mm, Waters, Germany) developing a two-step solvent gradient ranging from 2 to 20 % over 45 min and 20 to 32 % ACN in 0.08 % formic acid within 15 min, at a flow rate of 250 nL/min.

Full MS data were acquired in a range of 375-1,200 m/z with a maximum AGC target of 3e6 and automatic inject time at 120,000 resolution. Top 10 multiply charged precursors (2-5) over a minimum intensity of 5e3 were isolated using a 2 Th isolation window. MS/MS scans were acquired at a resolution of 60,000 at a fixed first mass of 110 m/z with a maximum AGC target of 1e5 or injection time of 118 ms. Previously isolated precursors were subsequently excluded from fragmentation with a dynamic exclusion of 120 seconds. TMT10 precursors were fragmented at a normalized collision energy (NCE) of 34 and TMTpro at a NCE of 32.

### Data analysis

Peptide identification was performed using CHIMERYS within the Proteome Discoverer 3.0 environment against the human reference proteome sequence database (UniProt; version: 2021-06-30). Searches were performed for specific tryptic peptides in the range of 7-30 amino acids with maximum 2 missed cleavages and maximum 3 modifications per peptide. Fragment mass tolerance was set to 20 ppm with maximum 15 candidates per MS/MS spectrum. Oxidation at methionine was set as variable modifications and the respective TMT reagents were selected as fixed modification. Peptide spectrum match (PSM), peptide and protein groups were filtered with a false discovery rate (FDR) of 1% using Percolator. S/N and intensity of RIs were extracted using the in-house developed Hyperplex (freely available: pd-nodes.org) at 10 ppm and assigned to the top hit per MS/MS spectrum for quantification. Post-processing was performed in the R environment if not indicated otherwise. For quantification PSMs were summed to peptides and protein groups. Single cell RI intensities are normalized to their sample loading within each analytical run. For HeLa/HEK and cardioid clustering, the raw RI intensities were log2 transformed, protein groups with less than 70% missing data across the entire dataset were imputed with random values from a normal distribution shifted into the noise. The RI intensities were then quantile normalized, batch corrected using ComBat for the analytical run and the TMT channel using the Perseus interface.^32^ Venn Diagrams are based on unique peptide sequences and are calculated using BioVenn.^33^ GRAVY scores were calculated for every unique peptide sequence identified from the respective condition, based on the Amino Acid Hydropathy Scores.^34^

## Data availability

All mass spectrometry-based proteomics data have been deposited to the ProteomeXchange Consortium via the PRIDE partner repository with the dataset identifier [PXD025387].

## Acknowledgements

We thank all members of our laboratories for helpful discussions. We specifically want to thank Clara Schmid for sharing cardioids with us and Elisabeth Roitinger for critical input on the manuscript. This work has been supported by EPIC-XS, project number 823839, the Horizon 2020 program of the European Union and the Vienna Science and Technology fund, project number LS20-079.

## Author contributions

D.H. and C.C. prepared and acquired samples. A.S. conceptualized and designed the proteoCHIP. C.C. performed data analysis and wrote the manuscript. G.T., S.M. and K.M. supervised the research and revised the manuscript.

