## Supplemental Figures for "An automated workflow for multiplexed single-cell proteomics sample preparation at unprecedented sensitivity"

### Title

### Affiliation

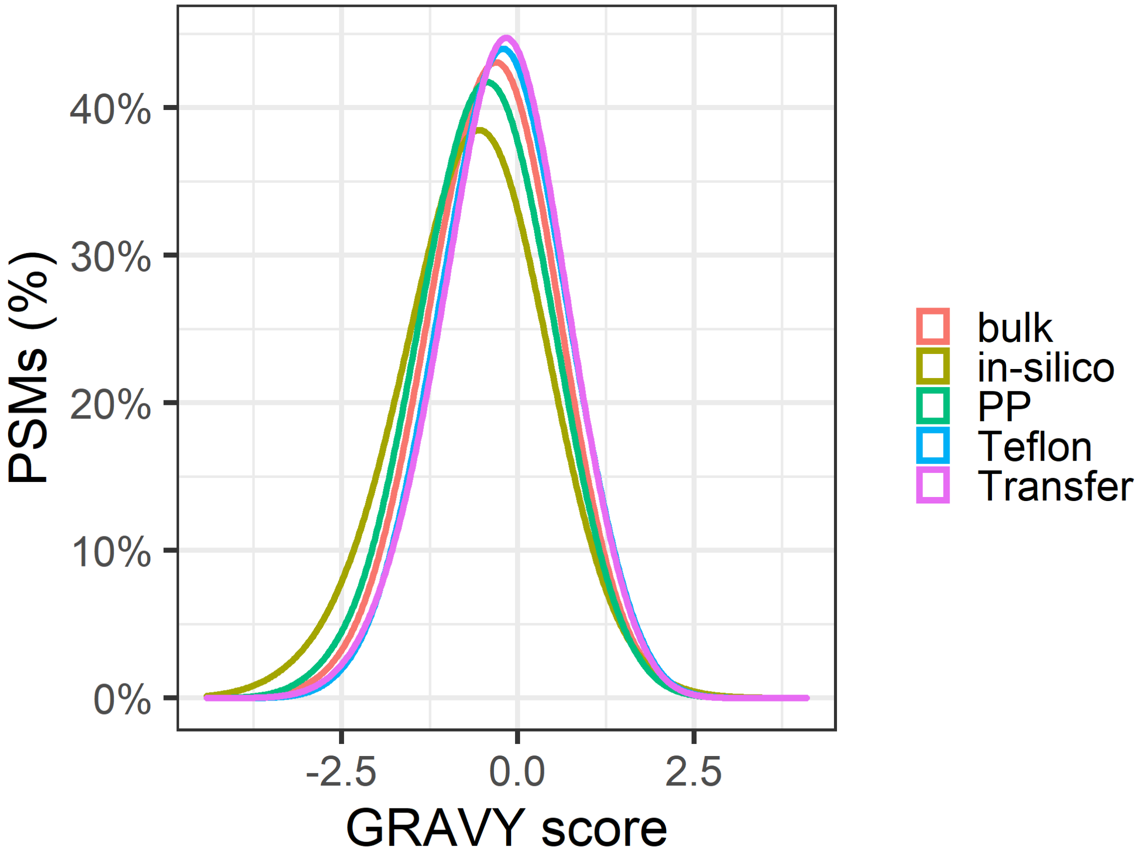


**Supplemental Figure 1: Surface comparison**. Gravy index of hydropathy of bulk HeLa digest in glass vials (bulk), in-silico digested human FASTA (in-silico), HeLa cells prepared standard plastic ware (PP), the proteoCHIP (Teflon) and prepared in the proteoCHIP but transferred to a standard PCR vial for injection (Transfer) across all PSMs.

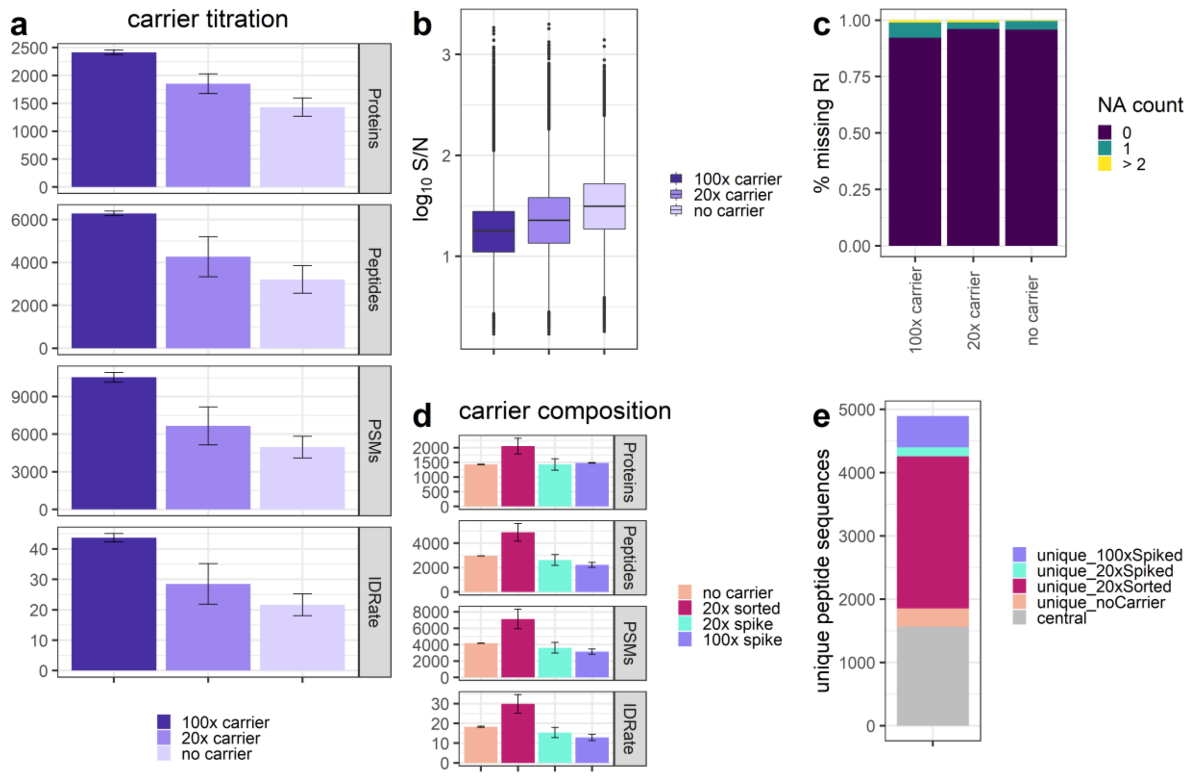


**Supplemental Figure 2: Carrier ratio influence on data quality. (a)** Protein groups, peptides, PSMs, MS/MS scans, ID-rate and **(b)** RI S/N and **(c)** missing quantitative data across all PSMs of single-cells with 100x, 20x sorted and no-carrier. **(d)** Protein groups, PSMs and **(e)** unique peptide sequence intersection for single-cells without carrier, with a 20x sorted carrier or 20x and 100x bulk spiked carrier.

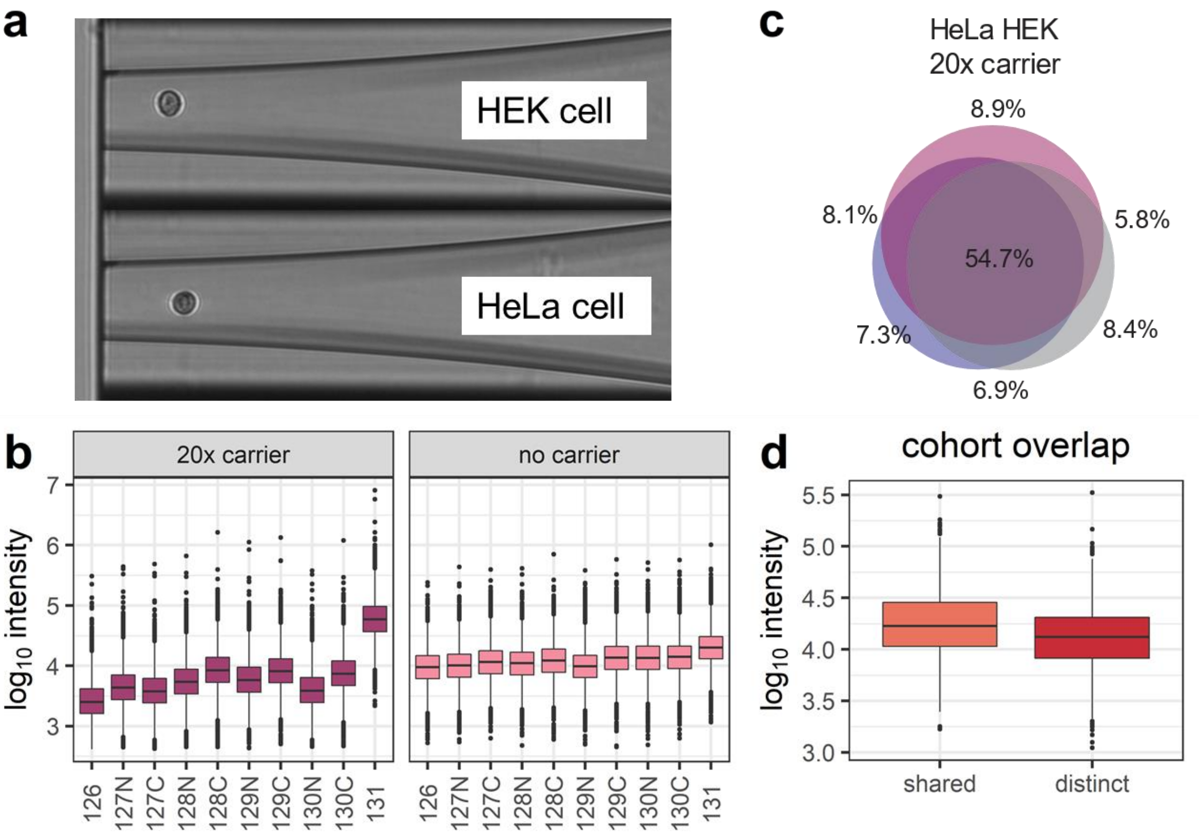


**Supplemental Figure 3: HeLa and HEK-293 cell type characterization. (a)** Image of a HEK-293 and a HeLa cell during image-based cell sorting using the cellenONE®. **(b)** RI intensity distribution across all channels for 20x and no-carrier samples. **(c)** Unique peptide sequence overlap of three HeLa/HEK-293 samples with 20x carrier. **(d)** $Log_{10}$ intensity overlap of shared or distinct unique peptides between 50 and 170 single-cells after filtering for 70% data completeness within each dataset.

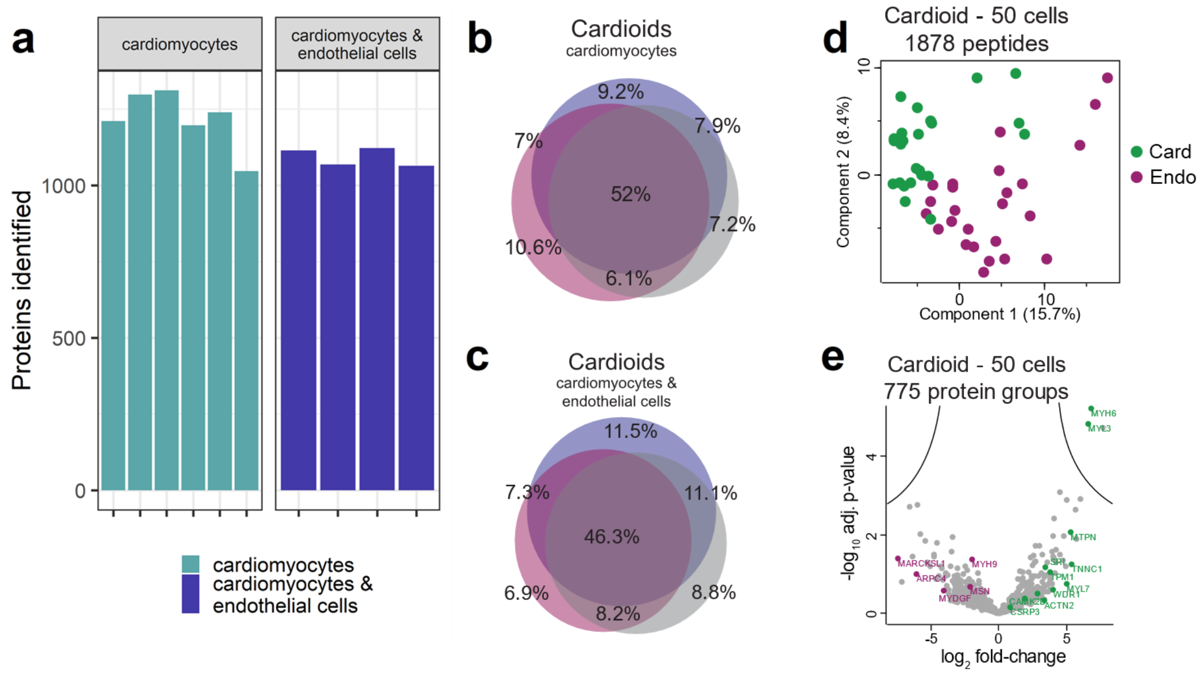


**Supplemental Figure 4: Cardioid sample cohort aggregation. (a)** Protein groups and **(b-c)** unique peptide sequences across three replicates with cardiomyocytes only or dual cardiomyocytes and endothelial cells. Single cell proteome **(d)** PCA and **(e)** cell type specific differences between cardiomyocytes (card - green) and endothelial cells (endo - purple), respectively. For volcano plots $\log_2$ fold change and $-\log_{10}$ adjusted p-value is shown.
